## Supplementary for "Forecasting cell fate during antibiotic exposure using stochastic gene expression"

**Supplementary Information**

| Construct | Primers |
|---|---|
| P$_{purA}$ | F: TTTATACAGTTCATCCATGCCCAGG<br>R: CCTGGGCATGGATGAACTGTATAA |
| P$_{inaA}$ | F: CAATGCTTTTCAGCGTTAACTCTG<br>R: ACGACAATGACTATAGGTGGT |
| P$_{rob}$ | F: CAAAATCTCAATACTTTTATTTCCG<br>R: TAATTGGATAATAGCATTTTTTGC |
| P$_{gadX}$ | F: GTTGACTACCTGGGTGGTC<br>R: ACTGTTTATTAATGTAGCACGCC |
| P$_{hdeA}$ | F: CCCCTGCTATCAATCTATGC<br>R: CACTGAGGTTATAACCTGGTTTTC |
| P$_{crp}$ | F: ACAGAGTACGCGTACTAACCAAATC<br>R: GGGCTATCAACTGTACTGCAC |
| P$_{\sigma70}$ | F: AATAATTCTTGACATTTATGCTTCCG<br>R: AATTGCACGTATTATACGAGC |
| P$_{rrnb\ \mathrm{P1}}$ | F: CTGAACAATTATTGCCCGTTTTACAG<br>R: GAATTAAACTTCGTAATGAATTACGTGTTC |
| P$_{acrAB}$ | F: GCCAGTAGATTGCACCGCG<br>R: TCGTGCTATGGTACATACATTCACA |
| P$_{sodA}$ | F: GGCAATCACGGCATTAAG<br>R: TTGGTTCATTATAGTTAATTAAATG |
| P$_{soxS}$ | F: CACGTTTATATCGCCGCTGATTG<br>R: TGTTGGGGAGTATAATTCCTCAAG |
| P$_{codB}$ | F: GATAATTTTTCCCCCACCTTTTTGC<br>R: CCGCCGCATTCTATTCATCTG |
| P$_{ompF}$ | F: CAAGTTATCTGTTTGTTAAGTCAAGCAATC<br>R: TGCAGGCATCTTTCCATTCAAAC |
| P$_{dnaQ}$ | F: GTTATGGATCCACTGGGTGATAC<br>R: CGCTATTTTACGCTATCGCGG |
| P$_{fis}$ | F: GCGAAGTGCGAGCAAGC<br>R: AGTTAAGAAATGACCATACTGTGACTGC |

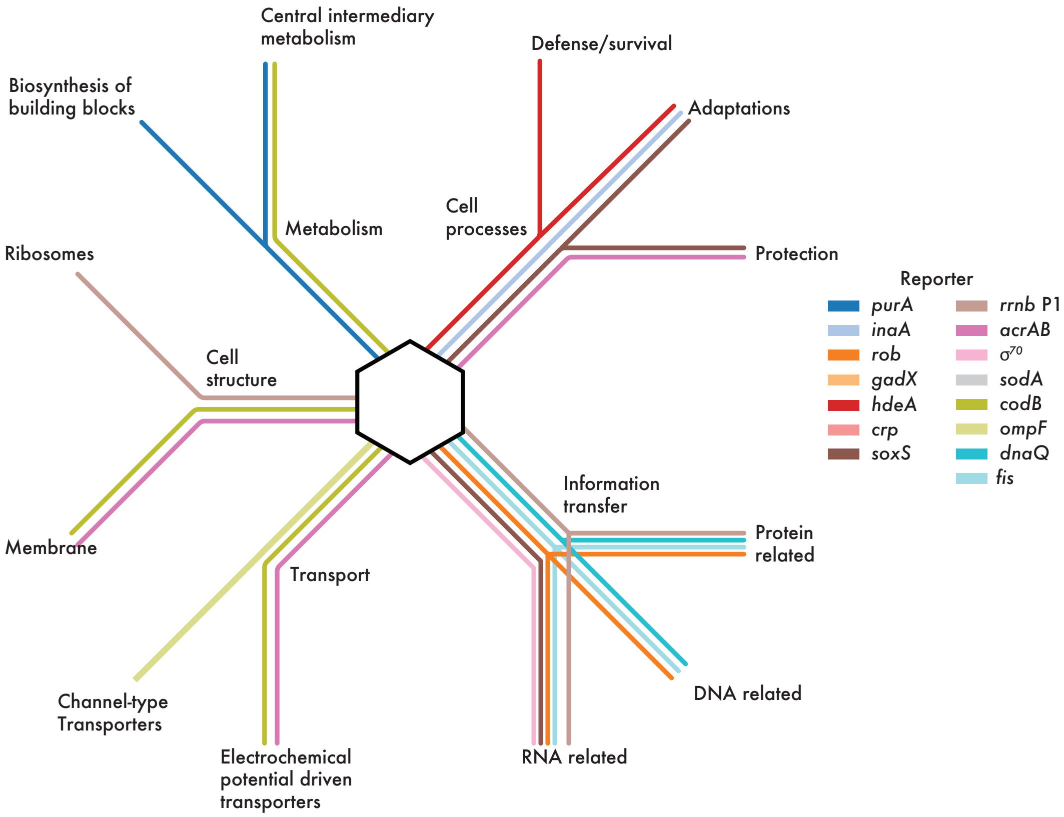


**Figure S1: Genetic ontology map for the promoters selected for study.**
The genetic ontology of each promoter is represented as a line connecting it through to each of its functional roles in the cell (Methods).

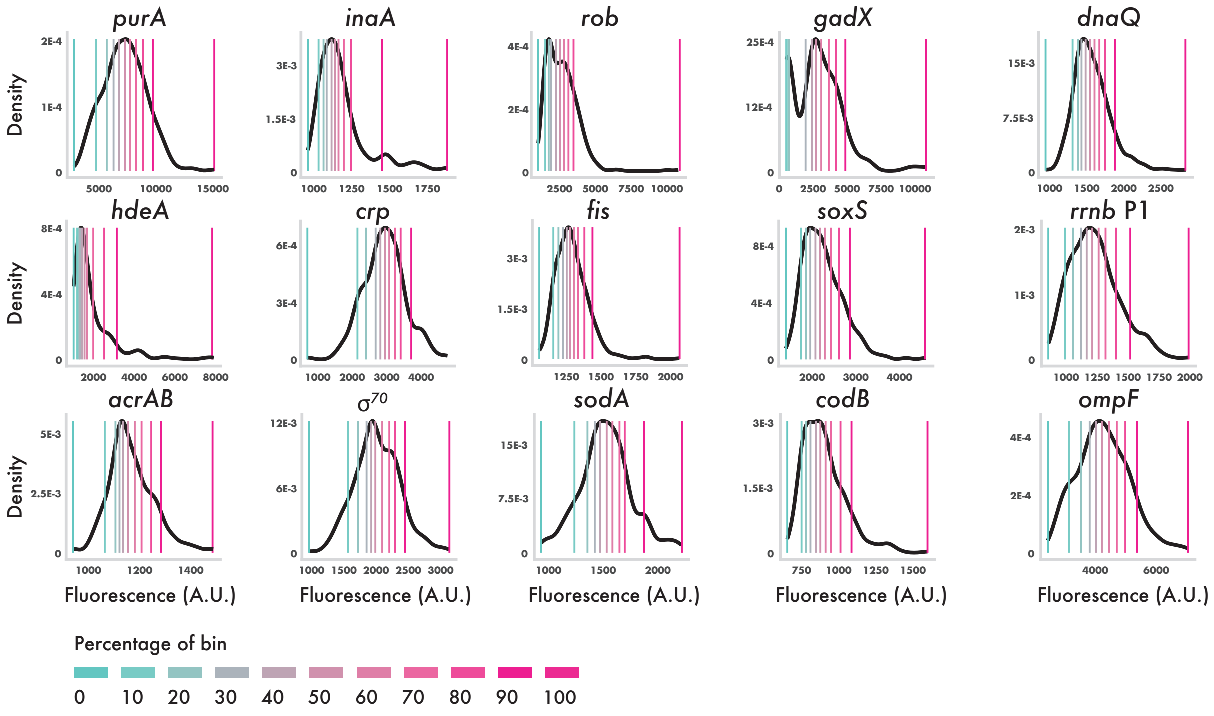


**Figure S2: Distributions and decile bins for all 15 promoters tested.**
Gray lines show histograms of the distributions of initial fluorescence at t = 0 for each of the 15 promoters. The colored lines show where each of the decile bins fall.

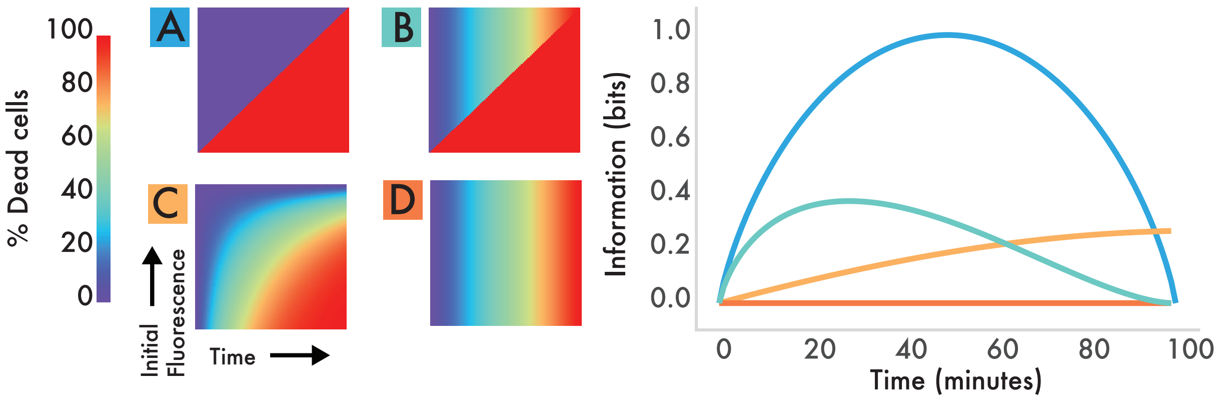


**Figure S3: Schematics of heatmaps showing variable information over time.**
Cartoon heatmaps of possible experimental patterns. **A** demonstrates a perfect split between alive and dead cells; the theoretical maximum information (1.0 bits) is achieved at the time of 50% cell death. **B** and **C** represent variations on this pattern. **D** demonstrates a heatmap schematic with the minimum information (0.0 bits) contained in initial fluorescence. Plots in the right panel show information over time for each of the heatmaps.

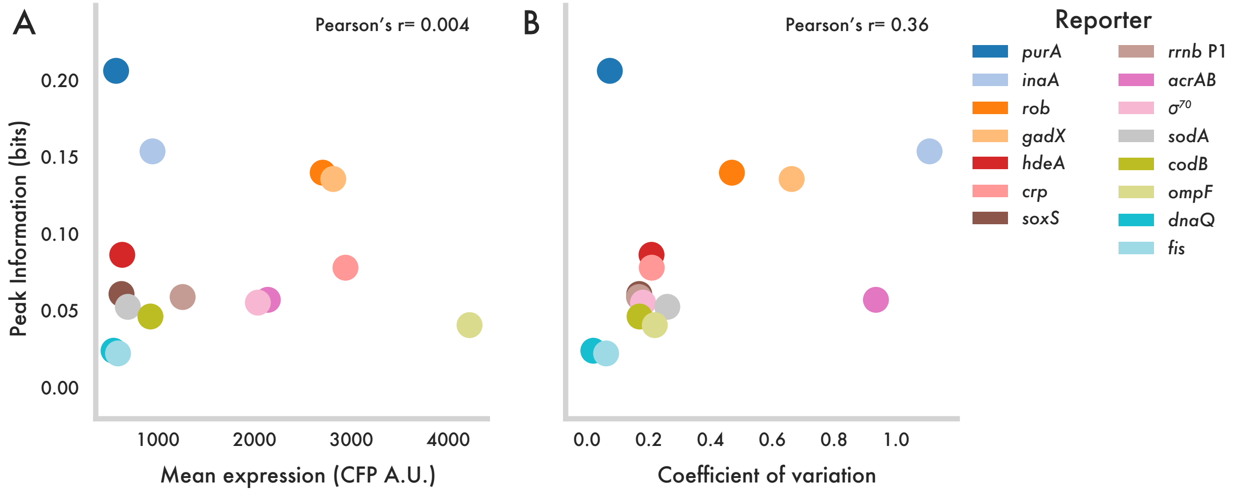


**Figure S4: Relationships between reporter statistics and information.**

(A) Scatter plot for mean expression versus peak information from Fig. 2B. Mean expression data taken from microscopy snapshots of each strain under the same CFP exposure time to ensure comparable statistics. Pearson's r-value calculates linear correlation between the two values (1.0 = positively correlated, 0.0 = uncorrelated).

(B) Scatter plot of coefficient of variation versus peak information for each of the promoters.

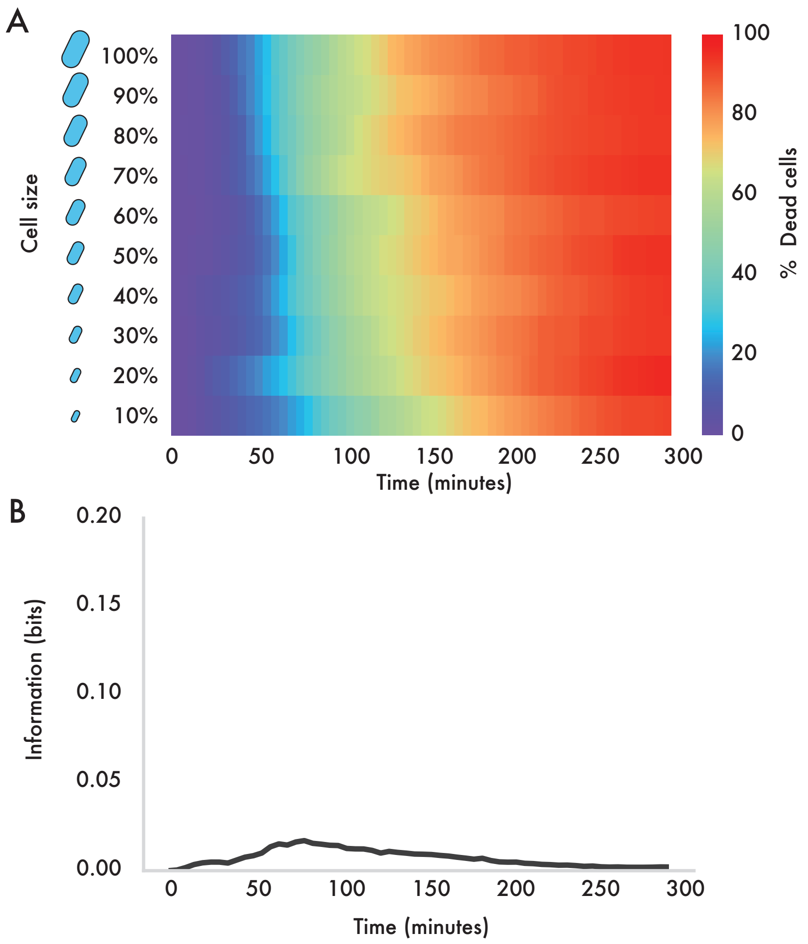


**Figure S5: Cell size as a predictor of cell fate.**

(A) Heatmap that mirrors Fig. 1C but uses initial cell size instead of fluorescence on the y-axis. Data are pooled from all carbenicillin reporter experiments.

(B) Information over time for cell size.

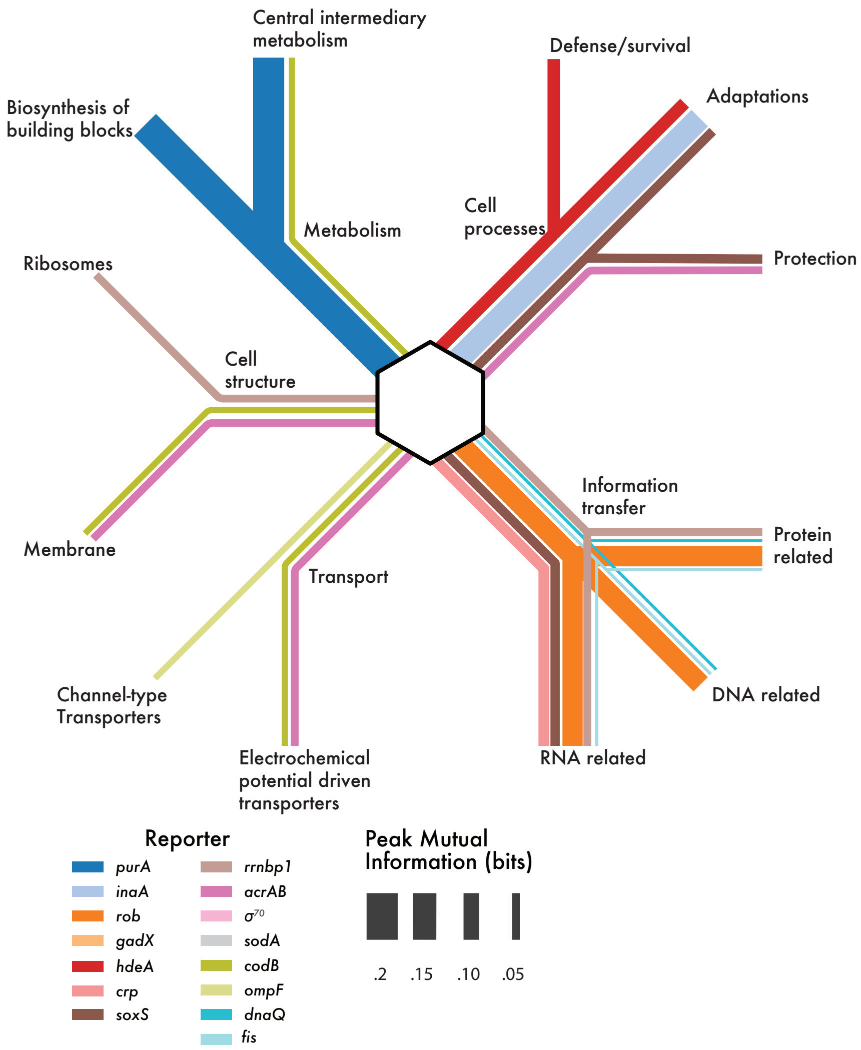


**Figure S6: Peak information from carbenicillin experiments mapped onto gene ontology.** The gene ontology map from Fig. S1 with the thickness of each connection dependent on the peak information from the carbenicillin data in Fig. 2B.

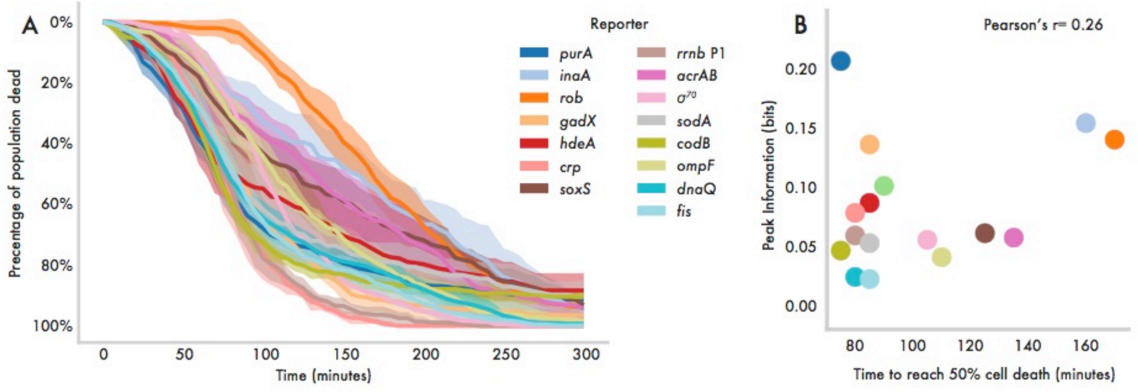


**Figure S7: Killing curves for all reporter strains over time under carbenicillin.**

(A) Killing curves for each strain extracted from time-lapse microscopy data. Solid lines represent the mean for at least 5 replicates while the shaded region represents the standard deviation across replicates.

(B) Scatter plot of time to reach 50% cell death versus peak mutual information. Pearson's r-value calculates linear correlation.

**Movie S1: Single-cell death under carbenicillin exposure for $P_{gadX}$-*cfp*.**

(A) Cell death over time. Red dot indicates particular point in time.

(B) Example microscopy movie of cells expressing $P_{gadX}$-*cfp* reporter (cyan) under carbenicillin exposure. Cell death is indicated by propidium iodide staining (red).

(C) Heatmap of cell death over time as a function of fluorescence at t = 0. White bar indicates particular point in time.

(D) Image showing fluorescence of individual cells at t = 0. As cells die they are eliminated from the image. Note that brighter cells tend to survive longer than dim cells.
